# Diverse ancestral representation improves genetic intolerance metrics

**DOI:** 10.1101/2024.11.04.621955

**Authors:** Alexander L. Han, Chloe F. Sands, Dorota Matelska, Jessica C. Butts, Vida Ravanmehr, Fengyuan Hu, Esmeralda Villavicencio Gonzalez, Nicholas Katsanis, Carlos D. Bustamante, Quanli Wang, Slavé Petrovski, Dimitrios Vitsios, Ryan S. Dhindsa

**Affiliations:** Department of Pathology and Immunology, Baylor College of Medicine, Houston, Texas 77030, USA; Jan and Dan Duncan Neurological Research Institute, Texas Children’s Hospital, Houston, TX 77030, USA; Centre for Genomics Research, Discovery Sciences, BioPharmaceuticals R&D, AstraZeneca, Cambridge, UK; Department of Bioengineering, George R. Brown School of Engineering, Rice University, Houston, TX, 77005, USA; Rice Neuroengineering Initiative, George R. Brown School of Engineering, Rice University, Houston, TX, 77030, USA; Department of Molecular and Human Genetics, Baylor College of Medicine, Houston, TX 77030, USA; Galatea Bio, Inc. Miami, FL 33016; Centre for Genomics Research, Discovery Sciences, BioPharmaceuticals R&D, AstraZeneca, Gaithersburg, US; Department of Medicine, Austin Health, University of Melbourne, Melbourne, Australia

## Abstract

Rapidly expanding genomic databases have enabled the identification of regions in the human genome intolerant to variation and thus likely relevant to human disease. However, despite their unprecedented scale, these datasets remain constrained by limited ancestral diversity. Here, we systematically evaluate how genetic diversity impacts gene- and sub-genic intolerance metrics by analyzing whole-exome sequencing data from 460,551 UK Biobank participants and 125,748 gnomAD participants across diverse ancestral backgrounds. Through comprehensive analysis of randomly sampled datasets with varying ancestral compositions, we demonstrate that genetic diversity, rather than sample size alone, drives the performance of intolerance metrics in identifying functionally critical genes and genic sub-regions. Notably, scores trained on variation observed in African and Admixed American ancestral groups showed superior resolution in detecting neurodevelopmental disease risk genes and haploinsufficient genes compared to scores trained on variation observed in European ancestry cohorts. Most strikingly, the Missense Tolerance Ratio (MTR) trained on 43,000 multi-ancestry exomes demonstrated greater predictive power than when trained on a nearly 10-fold larger dataset of 440,000 non-Finnish European exomes, indicating that European ancestry-based scores are approaching saturation. These findings establish ancestral diversity as the fundamental determinant for advancing intolerance metrics and underscore the urgent scientific and ethical imperative for enhanced population representation in genomic resources to fully realize the potential of precision medicine and drug discovery.

## Introduction

The emergence of population-scale sequencing datasets has enabled the identification of regions of the human genome that are intolerant to functional variation due to negative selection^1–5^. Broadly, intolerance scores provide a quantitative measure of the depletion of variants in a gene compared to a null expectation in the general population. We and others have developed intolerance metrics that quantify the intolerance of genes, genic sub-regions, and non-coding regions^2,3,6–12^. These scores have become a cornerstone in prioritizing genetic variants in diagnostic settings, discovering genes and genomic regions underlying human traits, and predicting drug targets^13^.

Gene-level intolerance metrics provide a valuable resource for studying the functional significance of human genes. For example, we previously introduced the Residual Variance Intolerance Score (RVIS), which uses standing variation in the human population to rank genes based on their tolerance to common functional variation (i.e., missense and protein-truncating variants)^1^. Other gene-level scores focus on more specific classes of mutations, such as missense Z (missense intolerance) and LOF-FDR and LOEUF (LOF intolerance)^4,5,14^. While gene-level metrics have proven extremely useful, certain regions of a gene can be more intolerant than other regions within the same gene, motivating the development of intragenic intolerance metrics. One example is the missense tolerance ratio (MTR), which identifies genic sub-regions that are intolerant to missense variation using a sliding window^9^.

As population-level sequencing datasets continue to expand in sample size, the genetic variants observed in these datasets also increase in number. Such an increase in the number of observed variants should, in theory, improve the performance of intolerance metrics. However, one of the persistent concerns is that a significant portion of current datasets are still disproportionately enriched for individuals of Northern European ancestry. Given that genetic variants and their frequencies differ dramatically across different populations^15^, the underrepresentation of global ancestries in these datasets not only exacerbates health inequities^16,17^, but also likely limits the resolution of genic intolerance metrics. Here, we explored the effect of ancestral diversity on genic and sub-genic intolerance metrics using two of the largest population-level exome sequencing databases, the Genome Aggregation Database (gnomAD) and the UK Biobank (UKB).

## Results

### gnomAD and UKB Cohort Characteristics

We leveraged large collections of exome sequence data from gnomAD (v2.1) and the UKB to measure the impact of ancestral diversity on genic and sub-genic intolerance metrics. We used previously existing ancestral group classifications for both these resources^14,17^. Briefly, these classifications were derived using PCA (Principal Component Analysis) methods to identify clusters of genetic similarity, which were then labeled based on established reference panels (see Methods).

The gnomAD v2 dataset includes aggregated allele frequency data from 125,748 exomes, among which we analyzed ancestry group specific allele frequencies from African (n=8,128), South Asian (n=15,308), Latino (n=17,296), East Asian (n=9,197), Ashkenazi Jewish (n=5,040), Non-Finnish European (n=56,885), and Finnish populations (n=10,824). In the UKB, we analyzed exome data from 460,551 individuals. Of these, 437,812 (95.06%) participants were of Non-Finnish European ancestry, 8,701 (1.89%) were of African ancestry, 2,671 (0.58%) were of Ashkenazi Jewish ancestry, 9,217 (2.00%) were of South Asian ancestry, and 2,150 (0.47%) were of East Asian ancestry.

As expected, the African ancestry cohorts (AFR) exhibited the most genetic diversity, relative to the current reference genome (hg38), followed by the Admixed American (AMR) and South Asian (SAS) cohorts. This difference was particularly apparent for common (MAF>0.05%) functional variants **(Tables 1, 2)**. For example, in gnomAD there was a 1.8-fold enrichment of common missense variants in the AFR cohort (n=142,178 variants among 8,128 individuals) compared to only the NFE cohort (n=878,771 variants in 56,885 individuals). Similarly, there was a roughly 1.6-fold enrichment of common PTVs in the AFR subset versus the NFE subset (**Table 1, 2**). The Finnish European group had the fewest common missense variants and PTVs, in part attributable to the founding bottleneck event that occurred roughly 120 generations ago, which reduced the effective gene pool and thus set their genetic history apart from other European populations^18,19^.

**Table 1.** Tally of protein-coding variants in the gnomAD ancestral groups. The number of common (MAF > 0.05%) and rare variants in each major genetic ancestry group observed in gnomAD. PTV = protein-truncating variant. Ancestral group abbreviations: AFR = African, AMR = Admixed American, SAS = South Asian; EAS = East Asian; ASJ = Ashkenazi Jewish; NFE = Non-Finnish European; FIN = Finnish.

| Variant Type | AFR<br>(n=8,128) | AMR<br>(n=17,296) | SAS<br>(n=15,308) | EAS<br>(n=9,197) | ASJ<br>(n=5,040) | NFE<br>(n=56,885) | FIN<br>(n=10,824) |
| --- | --- | --- | --- | --- | --- | --- | --- |
| **Common Missense** | 142,178 | 108,553 | 106,399 | 93,592 | 81,987 | 78,771 | 68,239 |
| **Common PTVs** | 5,226 | 4,148 | 4,201 | 4,031 | 3,673 | 3,233 | 3,121 |
| **Common Synonymous** | 118,445 | 86,892 | 80,336 | 67,901 | 61,886 | 60,227 | 50,958 |
| **Rare Missense** | 668,563 | 976,975 | 1,073,240 | 636,117 | 99,755 | 2,445,493 | 209,114 |
| **Rare PTV** | 46,402 | 69,696 | 76,656 | 46,263 | 8,959 | 211,411 | 18,491 |
| **Rare Synonymous** | 355,412 | 507,127 | 554,321 | 326,547 | 49,233 | 1,173,398 | 99,887 |

**Table 2.** Tally of protein-coding variants in the UK Biobank ancestral groups. The number of common (MAF > 0.05%) and rare variants in each major genetic ancestry group observed in the UK Biobank. NFE population was divided into three subsets of increasing sample size to test saturation for common variants. PTV = protein-truncating variant.

| Variant Type | AFR<br>(n=8,701) | SAS<br>(n=9,217) | EAS<br>(n=2,150) | ASJ<br>(n=2,671) | NFE<br>(n=20k) | NFE<br>(n=43k) | NFE<br>(n=440k) |
| --- | --- | --- | --- | --- | --- | --- | --- |
| **Common Missense** | 150,816 | 112,862 | 99,032 | 88,248 | 78,198 | 78,736 | 78,272 |
| **Common PTVs** | 7,647 | 6,278 | 5,718 | 5,474 | 4,650 | 4,630 | 4,684 |
| **Common Synonymous** | 125,683 | 84,416 | 72,860 | 65,668 | 59,161 | 59,429 | 59,271 |
| **Rare Missense** | 859,599 | 854,077 | 288,419 | 58,609 | 1,175,279 | 1,859,809 | 6,023,520 |
| **Rare PTV** | 69,882 | 70,874 | 22,267 | 6,639 | 112,946 | 189,338 | 772,007 |
| **Rare Synonymous** | 456,784 | 445,392 | 155,012 | 29,249 | 576,402 | 886,465 | 2,633,791 |

At current sample sizes, it appears that we have saturated the number of common (MAF > 0.05%) functional variants observed in Non-Finnish Europeans. To demonstrate this, we created three subsets of NFE UKB participants: one with 20,000 individuals, one with 43,000, and one with 440,000 individuals. The number of common variants was stable across these three cohorts (**Table 2**). However, the number of rare variants increased proportionally with sample size, apparent in the NFE subsets as well as the other major ancestral groups (**Table 1, 2**). Collectively, these results demonstrate that enhancing diversity, in addition to sample size, is critical for better representing standing variation in the global human population.

### Deriving Ancestry Group Specific Residual Variation Intolerance Scores

Given the significant differences in the amount of functional variation observed between populations, we hypothesized that lack of genetic diversity could limit the resolution of intolerance metrics. We first investigated this by creating ancestry group-specific versions of RVIS (**Supplementary Tables 1**,**2**). As previously described, RVIS regresses the total number of common missense and PTVs on the total number of protein-coding variants in a gene^1^. The score is then defined as the studentized residual for each gene, with a negative score indicating that a gene is more intolerant to functional variation.

The use of total number of variants on the X-axis in RVIS serves as an empirical proxy for that gene’s mutability and its length. Here, given differences in sample sizes and the number of observed variants across ancestral groups, we replaced the X-axis with genic mutability estimated from trimer mutation rates (see Methods). Encouragingly, genic mutability and total number of variants observed per gene were strongly correlated (UKB RVIS: Pearson’s r = 0.92; gnomAD RVIS: Pearson’s r = 0.94) (**Supplementary Fig. 1A, B)**.

We next calculated ancestry group-specific versions of RVIS in both the gnomAD and UKB datasets, using mutability on the x-axis and common (MAF > 0.05%) functional variants on the y-axis (**Fig 1A, C**). To assess differences in performance of these scores, we compared their ability to discriminate between genes that have been reported to be associated with severe disease and the rest of the exome. Specifically, we compiled five gene sets: monoallelic developmental and epileptic encephalopathy (DEE) genes (n=94), monoallelic developmental delay (DD) genes (n=435), monoallelic autism spectrum disorder (ASD) genes (n=190), haploinsufficient genes (n=360), and mouse essential genes (n=2,454) **(Supplementary Table 3)**.

**Figure 1.**
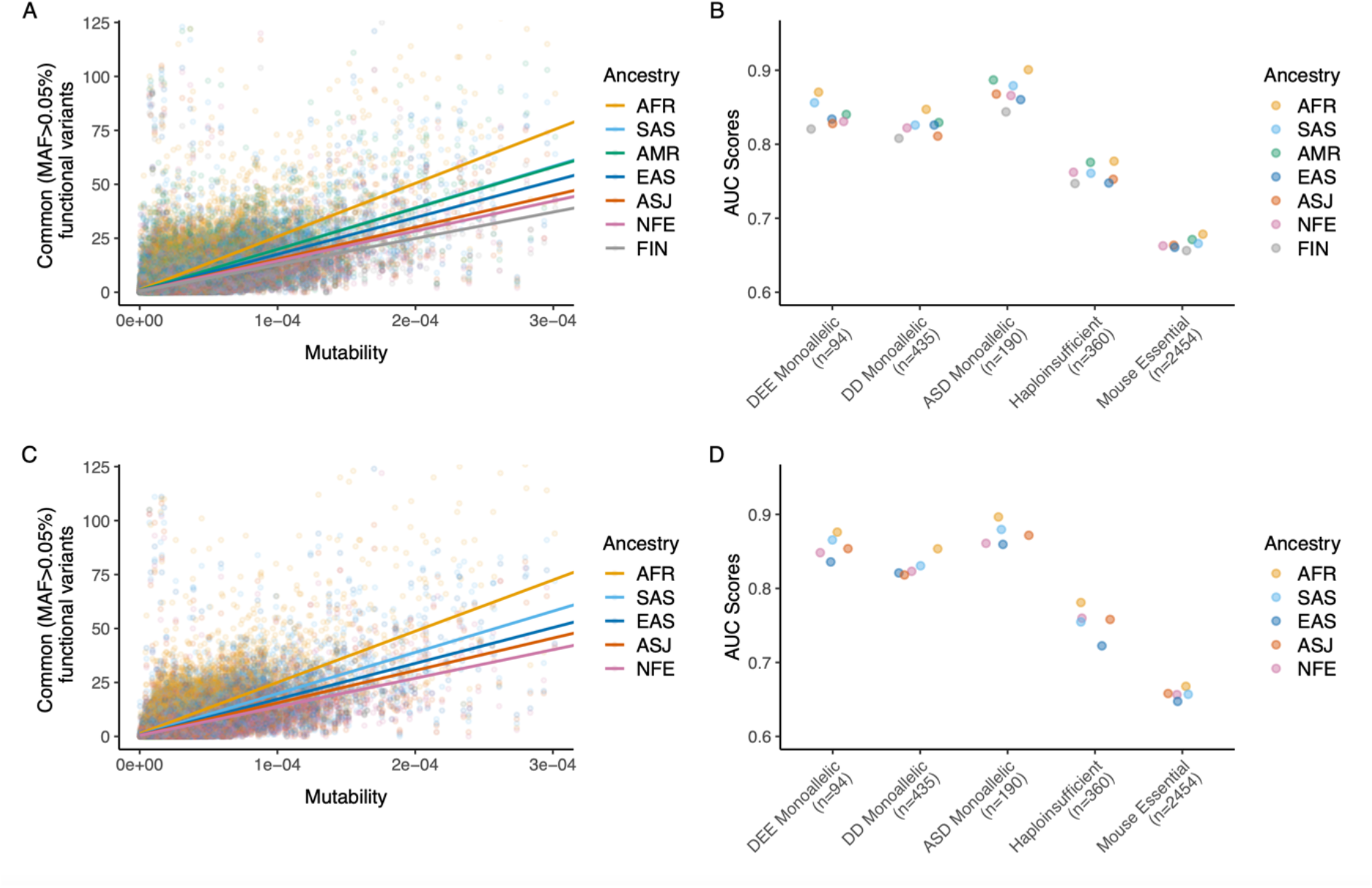
Ancestry Group Specific RVIS in gnomAD and UKB. **(A)** A scatter plot illustrating the regression of common (MAF > 0.05%) functional variants on mutability for seven different ancestry groups in gnomAD. The X- and Y-axes are capped to remove outlier genes. **(B)** AUC-ROC values of the ancestry group specific RVIS in gnomAD for predicting five gene lists. **(C)** Regression lines of common (MAF > 0.05%) functional variants versus mutability for five different ancestry groups in the UKB. **(D)** AUC-ROC values of the ancestry group specific RVIS in the UKB for predicting the following five gene lists. DEE = developmental and epileptic encephalopathy; DD = developmental delay; ASD = autism spectrum disorder.

Across all five gene sets, UKB- and gnomAD-derived RVIS scores that were trained on genetic variation data from the African ancestry cohort consistently achieved the highest area under the ROC curve (**Fig. 1B, D**). DeLong’s test demonstrated that RVIS_AFR_ AUCs were significantly higher than the RVIS_NFE_ AUCs for all gene sets excluding haploinsufficient genes (**Supplementary Table 4**,**5**). As expected, RVIS_AMR_ and RVIS_SAS_ also generally outperformed RVIS_NFE_. A sensitivity analysis demonstrated that this increased performance was robust to different MAF cutoffs for the Y-axis (**Supplementary Fig. 2**). Although the differences in AUC were modest, broad genetic representation in population-scale sequencing databases is clearly important in enhancing the resolution of RVIS.

### Increase in Gene-Level MTR Performance with Increasing Genetic Diversity

Whereas RVIS relies on the observation of common variants to quantify intolerance, other metrics are computed based on the total number of variants observed in a gene—irrespective of allele frequency—compared to a null expectation. Since the total number of observed variants correlates with sample size, we used the UKB to create down-sampled datasets of equal size but with varying compositions of ancestral diversity. Specifically, we defined one cohort of 42,739 exomes (“Maximally Diverse 43k”), which included data from individuals of African (n=8,701), South Asian (n=9,217), East Asian (n=2,150), and Ashkenazi Jewish (n=2,671) ancestry, as well as 20,000 randomly sampled Non-Finnish European samples **(Table 3)**. As a comparator, we defined a separate cohort of 42,740 NFE individuals (“NFE (n=43k)”). We also included two larger cohorts, including one with all 437,812 NFE samples (“NFE (n=440k)”) and a dataset that included all 460,551 of the above samples (“Full Dataset (n=460k)”). The “Maximally Diverse (n=43k)” cohort harbored 3,185,006 missense variants and PTVs compared to 2,132,513 in the “NFE only (n=43k)” cohort **(Table 3)**.

**Table 3.**
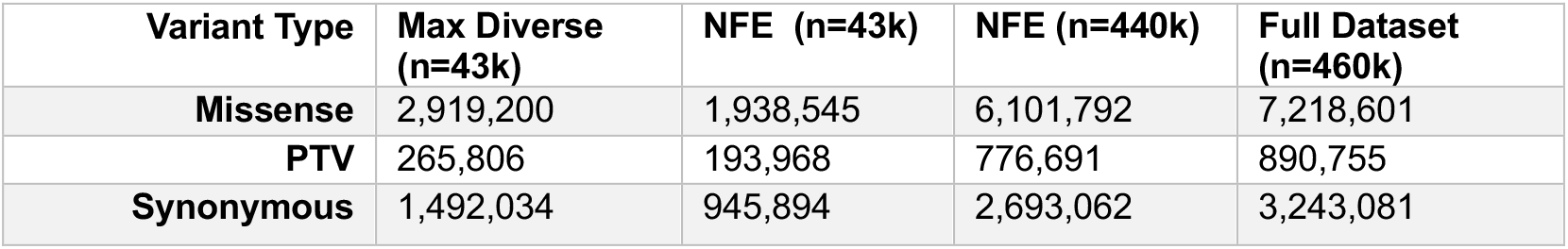
Total Number of Unique Variants and Variant Types. Total number of variants observed in each sub-sampled cohort from the UK Biobank as well as the full dataset. PTV = protein-truncating variants.

We then constructed gene-level missense tolerance ratio (MTR) scores using these four cohorts, as well as the ancestry-specific cohorts (**Supplementary Tables 6**)^9^. MTR is calculated by comparing the observed proportion of missense variants compared to an expected proportion given the sequence context of the gene (see Methods). Across all five gene sets, the “Maximally Diverse”-derived MTR scores achieved the best performance (**Fig. 2**). Interestingly, this score also outperformed the “Full Dataset”-derived score. One possibility is that the overrepresentation of NFE samples in the full dataset dilutes the signal gained from more genetically diverse cohorts. Furthermore, despite the drastic difference in sample size between the down-sampled (n=42,740) and entire (n=437,812) NFE cohorts, the performance of gene-level MTR in these subsets were generally comparable **(Supplementary Table 7)**.

**Figure 2.**
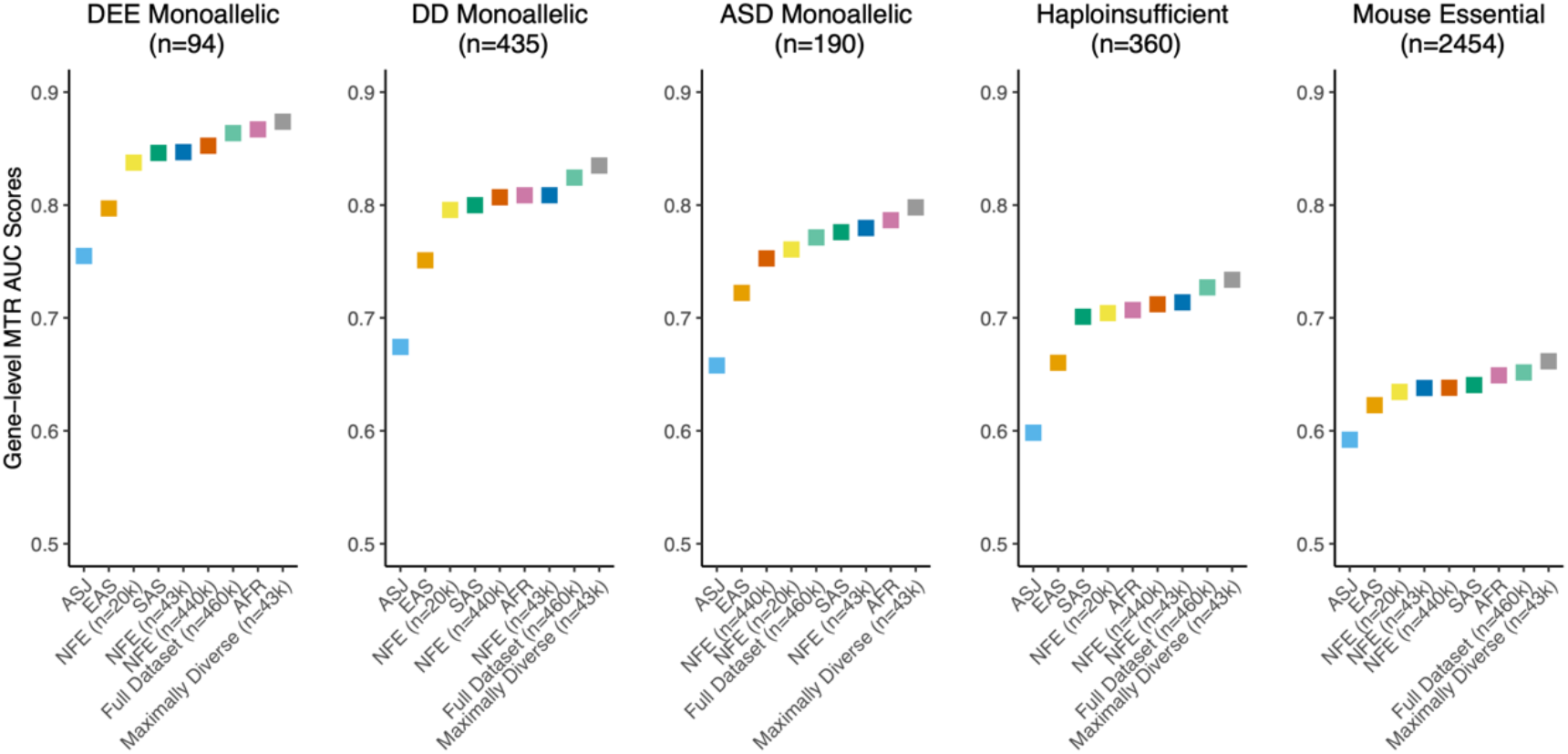
Performance of gene-level MTR scores. AUC-ROC scores illustrating the ability of each gene-level MTR score to predict five different gene lists. Each score represents a version of the score trained on the pre-defined UKB cohorts.

### Sliding Window MTR

We previously demonstrated that certain genic subregions can be under differential selection to missense variation using a sliding window version of MTR^7,9^. We thus next calculated a sliding window version of MTR for each UKB cohort, employing a 31-codon window (Methods). To compare the performance of these scores, we tested the ability of each to distinguish between ClinVar pathogenic/likely pathogenic variants and control variants. To define control variants, we included variants observed in either gnomAD or TopMED but not the UKB (**Fig. 3A, B**). We then compared the performance of these scores to predict *de novo* variants observed in affected probands versus those observed in their unaffected siblings from denovo-db^20^. AUCs were generally lower for this comparison, as not all *de novo* variants are necessarily pathogenic. However, this comparison revealed a similar pattern as the ClinVar analysis (**Fig. 3C**).

**Figure 3.**
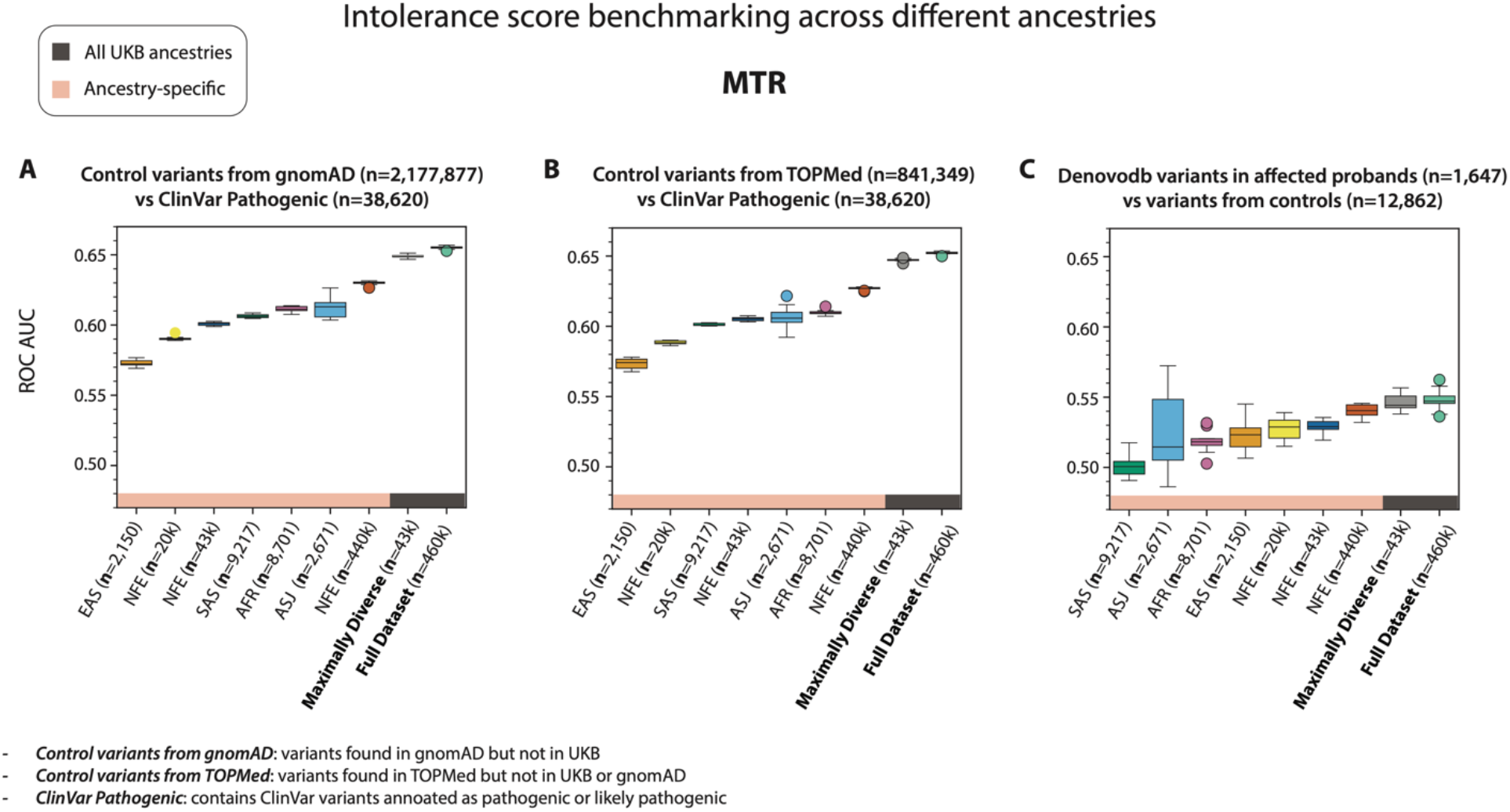
AUC values of sliding-window MTR in UKB for predicting three variant sets across different ancestry groups. **(A)** AUC-ROC scores reflecting each s0core’s ability to distinguish between ClinVar “pathogenic” or “likely pathogenic” variants and control variants from gnomAD. **(B)** Same as A, except control variants were derived from TopMED. **(C)** Ability of each score to distinguish between *de novo* variants observed in probands versus unaffected siblings in denovo-db.

Across all three variant sets, the versions of MTR trained on data from the most diverse cohorts achieved the best performance (i.e., the “maximally diverse cohort (n=43,000)” and the “full dataset (n=463,000)”). Strikingly, the performance of these two scores was nearly identical, even though the “maximally diverse” comprised of roughly 1/10^th^ the sample size. The remaining scores seemed to show a greater dependence on sample size than the gene-level metrics. This is likely because the region of interest is smaller for this version of the score (31 codon window versus an entire gene). Exemplifying this, the entire non-Finnish European cohort (n=440,000) outperformed each of the other individual ancestry groups for the three variant sets tested here.

### LOF O/E and LOF-FDR performance dependence with sample size

Finally, we examined the effects of genetic diversity on the performance of LOF intolerance metrics. Specifically, we considered our previously described LOF Observed/Expected metric (LOF O/E) and the related LOF-FDR score^4^ (**Supplementary Tables 8**,**9**). The LOF O/E score evaluates the ratio of LOF variants against the expected ratio of LOF variants under neutrality. The LOF-FDR score employs a one-sided binomial exact test with Benjamini and Hochberg false discovery rate multiple-testing correction to compare the observed to expected values (Methods). Scores trained with the entire NFE cohort (n=440k) and the complete dataset (n=460k) consistently outperformed the scores trained using reduced non-Finnish European cohort (n=42,740) and the maximally diverse cohort (n=42,739) across all gene sets (**Fig 4, Supplementary Fig. 3**). Compared to the prior metrics, sample size generally appeared to be the most important determinant of performance of LOF scores **(Supplementary Tables 10**,**11)**. The most plausible explanation for this observation is the relative scarcity of protein-truncating variants (PTVs) compared to missense and synonymous variants (**Table 1, 2**). PTVs are inherently rarer due to stronger purifying selection against deleterious mutations that disrupt gene function. As a result, even in ancestrally diverse populations, the absolute number of PTVs remains low, and thus larger sample sizes increase the confidence of observed statistics.

**Figure 4.**
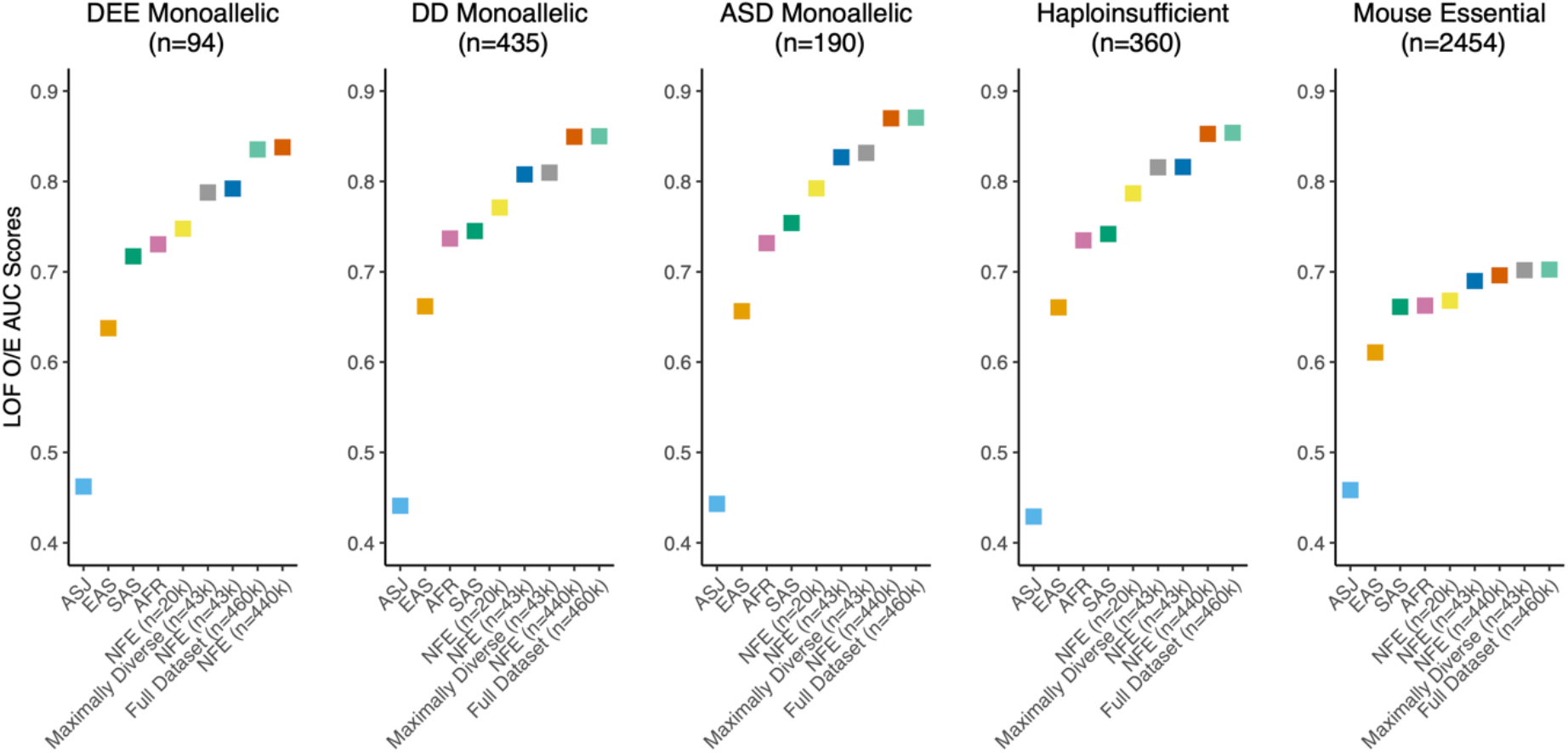
Performance of LOF intolerance metrics trained on different UKB cohorts. AUC-ROC scores illustrating the ability of LOF-OE to predict five different gene lists. Each score represents a version of the score trained on the UKB cohorts composed of different ancestries.

## Discussion

Improving our ability to discern intolerant regions of the human genome is crucial to prioritizing likely disease-causing mutations, facilitating the discovery of new risk genes, and enabling insights into human genome biology more broadly. Here, we performed an extensive study that evaluates the impact of ancestral diversity and sample size have on intolerance metrics using two of the largest population-level exome sequencing databases. Consistent with previous findings, we observed a higher prevalence of common functional variants in non-European ancestry groups compared to European populations. In turn, incorporating more diverse cohorts demonstrably improved the resolution of various intolerance metrics, which has also been previously observed non-coding intolerance metrics^10^. These differences were most apparent in RVIS and missense-specific intolerance scores (gene-level MTR and sliding window MTR). In comparison to RVIS and MTR, LOF intolerance metrics seemed more dependent on sample size. This is likely because LOF observations are considerably more sparse than missense and synonymous variants, meaning larger samples are required to achieve meaningful differentiation between test units. Altogether, our results broadly underscore the importance of achieving broad ancestral representation in our sequencing databases.

Our findings have broad implications beyond understanding evolutionary constraint on human protein coding genes. For example, the FDA’s recent guidance on Diversity Action Plans (2024) reinforces the importance of including underrepresented populations in clinical trials to improve the generalizability of study results. Our findings underscore this need by demonstrating that greater ancestral diversity in genetic studies improves intolerance metrics, which could lead to more equitable and effective precision medicine across populations. Secondly, increasing ancestral diversity in genetic datasets could have significant implications for drug discovery. A recent study demonstrated that genetically supported drug targets exhibit higher success rates in clinical trials^21^, further emphasizing the importance of incorporating diverse populations in the early stages of therapeutic development. Since the frequency of deleterious and protective alleles may vary across populations, broadening diversity in genomic data sets increases the community’s power to identify and derisk novel targets to move forward in drug development pipelines. Our work also has implications for understanding the transferability of results across populations. In related work, we and others^22–24^ have demonstrated that polygenic risk scores (PRS) models trained on European populations often perform poorly when applied to non-European groups, underscoring the importance of diversity in genetic research. The development of intolerance metrics that are more generalizable across different populations could potentially improve the portability of models that combine rare alleles and polygenic risk scores to model disease risk.

Finally, although we analyzed data from several ancestral groups, there still remains incomplete representation of many human populations. For example, the intolerance scores defined from AFR cohorts achieved the best performance, but we currently lack representation from majority of the African continent. In addition, the inclusion of diverse populations that have undergone strong demographic bottlenecks or high levels of consanguinity could also improve intolerance metrics^25^. Because there is an increased rate of homozygous LOF mutations in these populations, their increased representation would also enable the identification of genes intolerant to recessive variation, which has been notoriously difficult to assess with currently available data. As ongoing advancements in sequencing techniques lead to diminishing costs of exome and genome sequencing, it is imperative that this trend toward more diverse representation continue.

## Methods

### Genetic ancestry groups

Genetic ancestry group is distinct from race and ethnicity as explained by the National Academies of Sciences, Engineering, and Medicine (NASEM). While race and ethnicity are sociopolitical constructs based on perceived shared biological characteristics and culture, genetic ancestry refers to the specific paths through family trees where DNA is inherited from specific ancestors. Karczewski et al. describe the methodology gnomAD used to determine genetic ancestry group ^14^. We used the same ancestry labels for the UKB exomes as previously described^17,26^.

Splitting these exomes into geographic cohorts is beneficial in this study because genetic variants are observed with varying frequencies between genetic ancestry groups due to demographic histories, including bottleneck events. We want to underscore, however, that the genetic ancestry stratifications in both the UKB and gnomAD are artificially created and not naturally occurring. We recognize the concern behind genetic ancestry group annotations and their potential to exacerbate the illusion of natural ancestral populations as well as the association between geographic labels and socio-politically constructed concepts including race and ethnicity. Considering this and in accordance with recent guidelines from the National Academies of Sciences, Engineering, and Medicine (NASEM), we emphasize that genetic ancestry exists on a spectrum and relates to the family tree where DNA is inherited from specific ancestors. Moreover, genetic ancestry groups are stratified by genetic similarity, and the geographic labels from this study are artificially created.

### Estimating coverage-corrected gene-size and filtering qualifying variants

We used the gnomAD public transcripts as our coding-sequence source data (gnomAD Predicted Constraint Metrics v2.1.1, GRCh38). The data provided a canonical transcript per gene, defined as the longest isoform. Additional predicted constraint metrics including observed and expected variants counts per gene, observed/expected ratio, and Z-scores of the observed counts relative to expected were also included in the data. Mutability of missense, synonymous, and LoF variants per gene were also provided. The sum of these values was used as the mutability per gene. The UKB variants were annotated with SnpEFF^27^, and we filtered for the canonical transcript in the gnomAD.

Synonymous variants were identified with the function “synonymous”. Missense variants were labeled with the functions “missense”, “in frame insertion”, and “in frame deletion”. Loss of function variants were annotated with the functions “frameshift variant”, “stop gained”, “start lost”, “splice acceptor variant”, “splice donor variant”, “stop gained and frameshift variant”, “splice donor variant and coding sequence variant and intron variant”, “splice donor variant and intron variant”, “stop gained and in frame insertion”, “frameshift variant and stop lost”, “frameshift variant and start lost”, “start lost and splice region variant”, “stop gained and protein altering variant”, “stop gained and frameshift variant and splice region variant”, “stop gained and in frame deletion”, and “frameshift variant and stop retained variant”.

### Constructing RVIS

RVIS were calculated as described in Petrovski et al using a cohort of 125,748 exomes from the gnomAD (v2.1) and 460,551 exomes from the UKB^1^. Seven sets of ancestry group specific RVIS were computed from the gnomAD: AFR (n=8,128), SAS (n=15,308), AMR (n=17,296), EAS (n=9,197), ASJ (n=5,040), NFE (n=56,885), and FIN (n=10,824). Five sets of ancestry group specific RVIS were derived from the UKB Biobank (UKB): EAS (*n*=2,150), SAS (*n*=9,217), ASJ (*n*=2,671), AFR (*n*=8,701), and NFE (*n*=440k). The number of common functional variants per given gene were determined for each ancestry group. These common functional variants were defined as missense and loss-of-function variants with MAF values greater than 0.05%, which was a cutoff threshold validated by our sensitivity analysis (Supp 3). The number of common functional variants were then regressed against mutability, resulting in seven linear regression models for gnomAD and five models for the UKB. The studentized residuals from these models were computed as the ancestry group specific RVIS. Additionally, sensitivity analysis confirmed that the African ancestry group specific RVIS outperformed non-African ancestry group specific RVIS at most MAF thresholds (**Supplementary Fig. 2**).

### Constructing Gene-level MTR scores

We generated gene-level missense tolerance ratio (MTR) scores using the four cohorts and the ancestry-specific cohorts in the UKB. The formula for computing gene-level MTR was

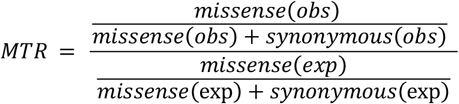

which compares the ratio of observed fraction of missense variants relative to observed missense and synonymous variants against the expected fraction of missense variants. With the assumption that synonymous variants are evolutionarily neutral and free from selective pressures, the MTR computes the fraction of allowed missense mutations relative to the maximum possible proportion of missense variation in the absence of selective pressure. The gene-level MTR focuses on the cumulative intolerance of the whole gene.

### LOF O/E and LOF-FDR

As previously described, we generated LOF O/E and LOF-FDR scores using the four cohorts and the ancestry-specific cohorts in the UKB^4^. Briefly, the expected rate is calculated by taking the mutation rate of LOFs (i.e., PTVs) and dividing that by the sum of mutation rates of all possible mutation effects in the gene, including synonymous, missense, and LOF variants. Mutability estimates were derived from the gnomAD v2.1 metrics file^14^. To account for frameshift variants, we multiplied the PTV mutability by 1.25 as previously described^5^.

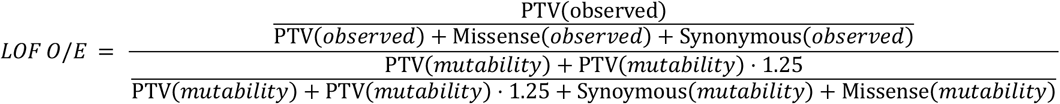

To calculate LOF-FDR, we computed a one-sided binomial exact test using the number of LOF variants, total number of unique variants, and expected fraction of LOF variants under neutrality and applied Benjamini & Hochberg false discovery rate multiple-testing correction.

### Benchmarking Gene-Level Scores

To test cohort specific gene-level scores in determining disease-causing genes, we performed simple logistic regression models for RVIS, gene-level MTR, LOF O/E, and LOF FDR (Supplementary Table 4,5,7,9,10). In these models, the gene-level metric was used as the predictor variable, while a binary variable distinguishing disease-causing genes from the rest of the HGNC genes served as the response variable. For each pairing of test set and gene-level score, the area under the ROC curve (AUC) for identifying disease causing genes was computed. DeLong’s test was used to assess for significant difference in the predictive ability between models.

Five gene sets were used as test sets to evaluate performance of the cohort specific gene-level scores. These were monoallelic developmental and epileptic encephalopathy (DEE) genes, monoallelic developmental delay (DD) genes, monoallelic autism spectrum disorder (ASD) genes^28^, haploinsufficient genes^29^, and mouse essential genes^30^. The monoallelic ASD gene list was curated by selecting SFARI Tier 1 ASD genes (n=207) and filtering genes with biallelic mechanism, leaving 190 total monoallelic ASD genes. The monoallelic DD gene list was determined by selecting genes with “Brain/Cognition” and “Definitive” labels from the Developmental Disorder Genotype-Phenotype Database (DD2GP) then filtering for monoallelic genes (n=218) and combining these genes with 199 genes found to be significant from the Kaplanis et al, culminating in total of 435 genes for the DD monoallelic set^31^. The monoallelic DEE gene list was created by selecting for “Developmental and Epileptic Encephalopathy” phenotype genes from Online Mendelian Inheritance in Man (OMIM) and combining with the DEE genes from the Epi25k study of epilepsy ^32,33^. We provided additional information on curating monoallelic DEE, DD, and ASD gene lists in our previous study^28^.

### Sliding Window MTR variant-level benchmark

Sliding window MTR scores were calculated as described in Traynelis et al. and Vitsios et al. for every codon position of 18,823 canonical transcripts over a sliding window of 31 codons^7,9^. We employed variant-level data from nine different cohorts (of single or diverse ancestry groups) derived from the UK Biobank (UKB): EAS (*n*=2,150), SAS (*n*=9,217), ASJ (*n*=2,671), AFR (*n*=8,701), NFE (*n*=20k), NFE (*n*=43k), NFE (*n*=440k), all-ancestry groups (*n*=43k) and all-ancestry groups (*n*=460k). To test MTR’s ability to discriminate pathogenic missense variants from benign, we composed three sets of benign variants and two sets of pathogenic variants. The benign sets we compiled comprise of: i) variants present in gnomAD (v2), but not present in UKB, ii) variants present in TOPMed, but not present in UKB or gnomAD (v2), and iii) de novo variants from denovo-db present in control individuals from the SSC or non-SSC samples^20^. As pathogenic variant sets, we used: i) variants confidently annotated in ClinVar as “Pathogenic” or “Likely pathogenic”, or ii) de novo variants from denovo-db present in affected individuals from the SSC or non-SSC samples.

To test MTR’s performance, as calculated across the different cohorts, in distinguishing pathogenic missense variants form benign, we used MTR as a measure of probability that a variant is benign: the higher the MTR score, the greater the probability. For each combination of test sets and MTR metrics, the area under the ROC curve (AUC) for distinguishing pathogenic from benign variants was calculated. Due to the imbalance in the respective positive and negative test sets, we randomly down-sampled the larger one in each case to match the size of the other set of the pair. Finally, we repeated the down-sampling and calculated AUC scores across 10 iterations for each pair of pathogenic-benign variant sets to include enough data points and increase the robustness of the discrimination tasks.

## Acknowledgments

We thank the participants in the UK Biobank and gnomAD for their contributions to research (UKB Resource Application Numbers 26041 and 65851). We extend our gratitude to the AstraZeneca Centre for Genomics Research Analytics and Informatics team for processing and analysis of sequencing data. R.S.D. is supported by NIH DP5 OD036131 and a Longevity Impetus Grant from Norn Group, Hevolution Foundation, and Rosenkranz Foundation.

## Data availability

Gene-level intolerance scores are included as supplementary tables and sliding-window MTR scores are available on FigShare: https://figshare.com/s/ac3dc3caf65af0162fad

All UKB whole-exome sequencing data described in this paper are publicly available to registered researchers through the UKB data access protocol. Exomes can be found in the UKB showcase portal: https://biobank.ndph.ox.ac.uk/showcase/label.cgi?id=170. Additional information about registration for access to the data is available at http://www.ukbiobank.ac.uk/register-apply/. Data for this study were obtained under Resource Application Number 26041 and 65851.

The following publicly available datasets and databases were used:

Gene lists: https://github.com/macarthur-lab/gene_lists

gnomAD v2.1.1: https://gnomad.broadinstitute.org/

denovo-db v1.6.1: https://denovo-db.gs.washington.edu/denovo-db/

## Code availability

Code for generating gene-level scores is available at: https://github.com/alhanster/ancestral_diversity. The code for generating sliding-window MTR scores is available at: https://github.com/astrazeneca-cgr-publications/OncMTR/tree/mtr-ancestry-specific

## Ethics declarations

The protocols for the UK Biobank are overseen by The UK Biobank Ethics Advisory Committee (EAC); for more information see https://www.ukbiobank.ac.uk/ethics/ and https://www.ukbiobank.ac.uk/wp-content/uploads/2011/05/EGF20082.pdf.

