## Supplementary Information for "Diverse ancestral representation improves genetic intolerance metrics"

### Supplementary Figure 1

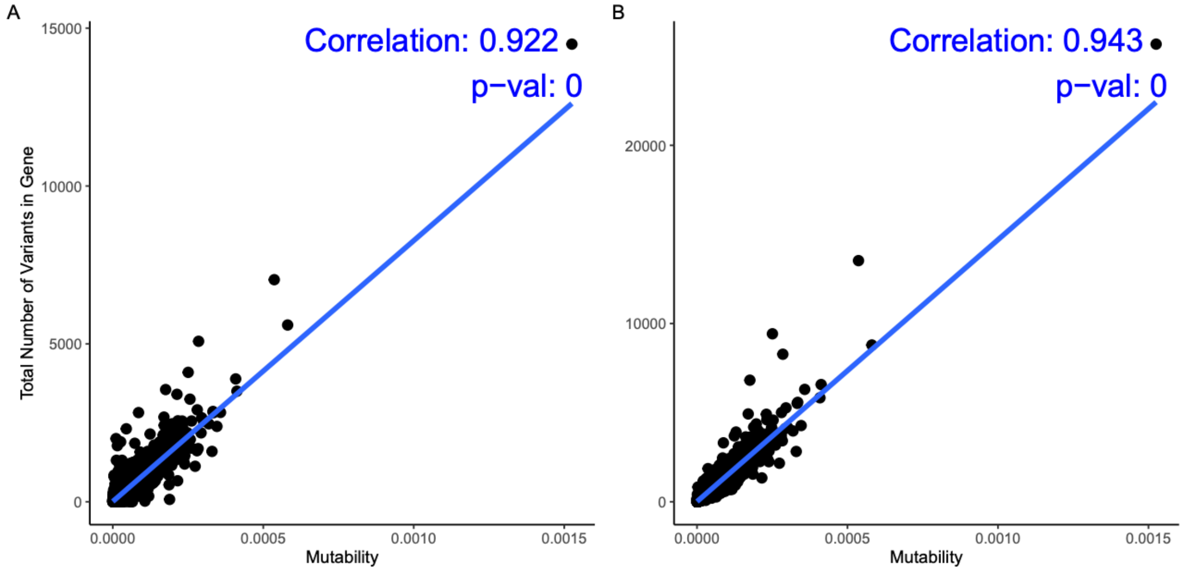

**Correlation of Mutability and total observed number of variants in gene for UKB [A] and gnomAD [B] cohorts.** Mutability values were compared with total number of observed variants for the given gene in UKB and gnomAD cohorts. Correlation calculated via Pearson's R.

### Supplementary Figure 2

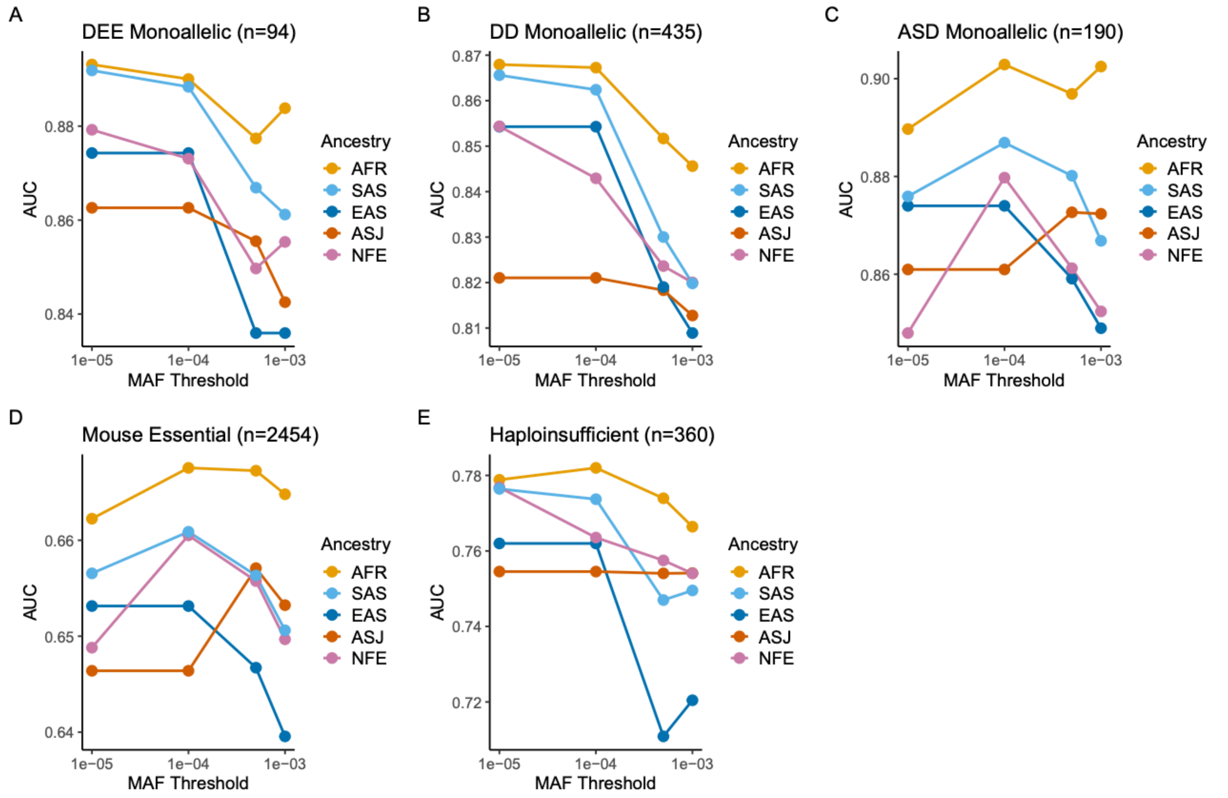

**Sensitivity Analysis to determine MAF cut-off.** RVIS scores were computed by varying MAF cut-offs. Performance of the RVIS scores were evaluated across different gene-lists. Sensitivity analysis validated our MAF cut-off value of 0.05%.

### Supplementary Figure 3

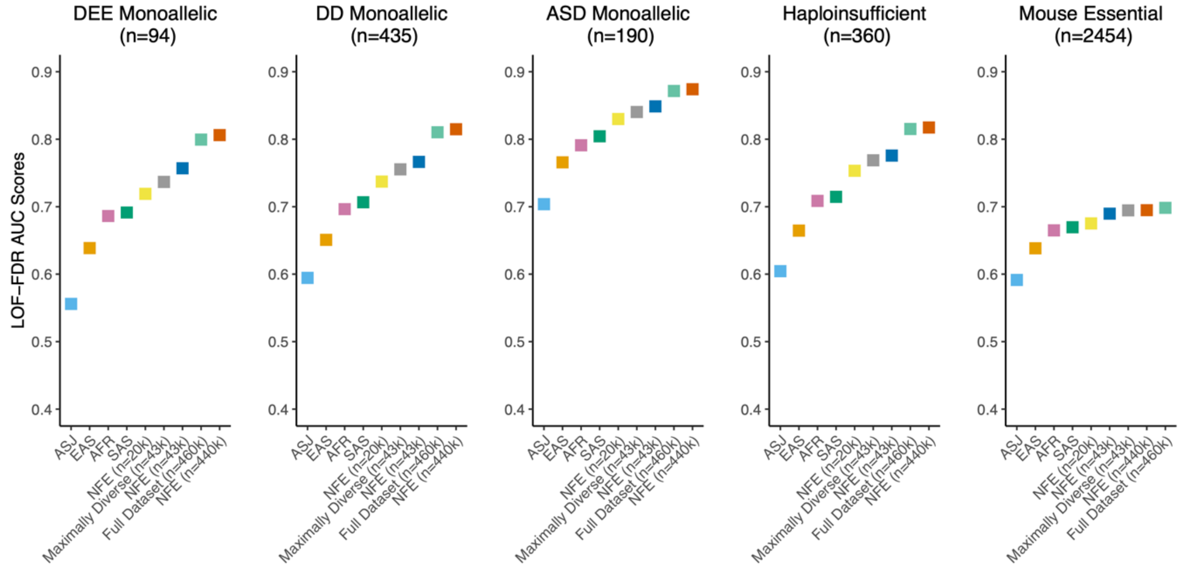

**Performance of LOF-FDR trained on different UKB cohorts.** AUC-ROC scores illustrating the ability of LOF-FDR to predict five different gene lists. Each score represents a version of the score trained on the UKB cohorts composed of different ancestries.

### Supplementary Table Legends

#### Supplementary Table 1 – UKB RVIS Scores

Provided as external file. Genes and UKB-derived RVIS scores are described in the table. Variable “x” accounts for total number of variants. Variable “y” indicates respective counts by ancestry (AFR = African, ASJ = Ashkenazi Jewish, EAS = East Asian, SAS = South Asian, NFE = non-Finnish European). Mutability values for each gene was derived from the gnomAD metrics table.

#### Supplementary Table 2 – gnomAD RVIS Scores

Provided as external file. Genes and gnomAD-derived RVIS scores are described in the table. Variable “x” accounts for total number of variants. Variable “y” indicates respective counts by ancestry (AFR = African, AMR = Admixed American, ASJ = Ashkenazi Jewish, EAS = East Asian, FIN = Finnish, NFE = Non-Finnish European, SAS = South Asian). Mutability values for each gene was derived from the gnomAD metrics table.

#### Supplementary Table 3 – Gene Lists

Provided as external files. Each gene-list is present on separate sheets. From left to right: DD Monoallelic, DEE Monoallelic, ASD Monoallelic, Clingen Haploinsufficiency, and Mouse Essential.

#### Supplementary Table 4 – Logistic regression and DeLong test for UKB-derived RVIS

Provided as external files. In the Logistic Regression sheet, ancestry-specific RVIS scores, gene-list, p-value, and corresponding AUC values provided (AFR = African, ASJ = Ashkenazi Jewish, EAS = East Asian, SAS = South Asian, NFE = non-Finnish European). In the DeLong Test sheet, gene-list, score1, score2, and p-value are provided.

#### Supplementary Table 5 – Logistic regression and DeLong test for gnomAD-derived RVIS

Provided as external files. In the Logistic Regression sheet, ancestry-specific RVIS scores, gene-list, and corresponding AUC values provided (AFR = African, AMR = Admixed American, ASJ = Ashkenazi Jewish, EAS = East Asian, FIN = Finnish, NFE = Non-Finnish European, SAS = South Asian). In the DeLong Test sheet, gene-list, score1, score2, and p-value are provided.

#### Supplementary Table 6 – UKB MTR Scores

Provided as external file. Genes and UKB-derived MTR scores are described in the table. Variables “mis” and “syn” indicate counts of missense and synonymous variants for the individual ancestry and compiled groups (AFR = African, ASJ = Ashkenazi Jewish, EAS = East Asian, SAS = South Asian, NFE = non-Finnish European, Maximally Diverse n=43k, NFE only n=20k, NFE only n=43k, NFE only n=440k, and Full Dataset n=460k). Variables “possible_mis” and possible_syn” account for expected counts of missense and synonymous variants.

#### Supplementary Table 7 – Logistic regression and DeLong test for UKB-derived MTR

Provided as external files. In the Logistic Regression sheet, cohort-specific MTR scores, gene-list, p-value, and corresponding AUC values provided (AFR = African, ASJ = Ashkenazi Jewish, EAS = East Asian, SAS = South Asian, NFE = non-Finnish European). In the DeLong Test sheet, gene-list, score1, score2, and p-value are provided.

#### Supplementary Table 8 – UKB LOF O/E

Provided as external file. Genes and UKB-derived LOF O/E scores are described in the table. Variable “lof” indicate respective counts of LOF variants for the individual ancestry and compiled groups (AFR = African, ASJ = Ashkenazi Jewish, EAS = East Asian, SAS = South Asian, NFE = non-Finnish European, Maximally Diverse n=43k, NFE only n=20k, NFE only n=43k, NFE only n=440k, and Full Dataset n=460k). Variable “total” indicate respective counts of total variants for the individual ancestry and compiled groups. Variable “possible_lof” accounts for expected counts of LOF variants. Variable “mu_lof” was derived from the gnomAD metrics table and accounts for mutability values for LOF variants in a gene. Variable “exp_lof_percent” was calculated using mutability values from the gnomAD metrics table (Methods).

#### Supplementary Table 9 – UKB LOF-FDR

Provided as external file. Genes and UKB-derived LOF-FDR scores are described in the table. Variable “lof” indicate respective counts of LOF variants for the individual ancestry and compiled groups (AFR = African, ASJ = Ashkenazi Jewish, EAS = East Asian, SAS = South Asian, NFE = non-Finnish European, Maximally Diverse n=43k, NFE only n=20k, NFE only n=43k, NFE only n=440k, and Full Dataset n=460k). Variable “total” indicate respective counts of total variants for the individual ancestry and compiled groups. Variable “possible_lof” accounts for expected counts of LOF variants. Variable “exp_lof_percent” was calculated using mutability values from the gnomAD metrics table (Methods).

#### Supplementary Table 10 – Logistic regression and DeLong test for UKB-derived LOF O/E

Provided as external files. In the Logistic Regression sheet, cohort-specific LOF O/E scores, gene-list, p-value, and corresponding AUC values provided (AFR = African, ASJ = Ashkenazi Jewish, EAS = East Asian, SAS = South Asian, NFE = non-Finnish European). In the DeLong Test sheet, gene-list, score1, score2, and p-value are provided.

#### Supplementary Table 11 – Logistic regression and DeLong test for UKB-derived LOF-FDR

Provided as external files. In the Logistic Regression sheet, cohort-specific LOF-FDR scores, gene-list, p-value, and corresponding AUC values provided (AFR = African, ASJ = Ashkenazi Jewish, EAS = East Asian, SAS = South Asian, NFE = non-Finnish European). In the DeLong Test sheet, gene-list, score1, score2, and p-value are provided.
